# Phylospatial optimal transport for single-cell and spatial transcriptomics data integration

**DOI:** 10.64898/2026.11.11.765961

**Authors:** Pedro F. Ferreira, Oskar Kviman, Jens Lagergren

## Abstract

The spatial organization of evolving cell populations in tumors influences how they progress and respond to treatment. Spatially resolved transcriptomics data may be used to examine the locations of these diverse cell states in tumor sections. However, these technologies often lack single-cell resolution and instead capture groups of cells in spatial spots, making it difficult to resolve cancer clones. In contrast, dissociated single-cell transcriptomics data are better suited for obtaining cell lineages, but do not capture the locations of the cells in the tissue. Here, we introduce SpOTr, an optimal transport approach that leverages the concordance between the evolutionary tree and the spatial distribution of cancer cells to integrate single-cell and spatial transcriptomics data from tumors. SpOTr combines a spot deconvolution model with Fused Gromov–Wasserstein optimal transport, in which cells that are close in the tree are drawn to nearby spots. In simulations and in real spatial lineage tracing data, SpOTr recovered the spatial locations of clones better than methods that ignore the tree. In a human colorectal liver metastasis, SpOTr mapped copy number clones from dissociated cells to spatial spots, showing that it successfully maps the evolutionary history of tumors in space.

## Introduction

Intra-tumor heterogeneity is a hallmark of cancer [1] and one of the main causes for treatment failure [2]. Cancer cells often acquire genomic aberrations as they divide, resulting in subclones of cells with distinct genotypes that may result in distinct cell states [3, 4]. On the one hand, the interactions between these cells and the surrounding tissue further contribute to the emergence of novel phenotypes and the formation of a complex tumor microenvironment (TME) [5]. On the other hand, the physical constraints imposed by the spatial structure of the tumor create growth patterns that shape its evolutionary dynamics and the localization patterns of subclones [6, 7]. However, the relationship between subclonal phenotypes and tumor evolution is still unclear [8, 9]. To address this question, the gene expression of cancer clones must be analyzed in the context of their spatial locations in the tumor.

Recent technological advances have enabled researchers to study intra-tumor heterogeneity in its spatial context through spatially-resolved transcriptomics (SRT) [10, 11]. These technologies quantify gene expression in two-dimensional sections of tumors, allowing insights into the spatial distribution of cancer cell states [12], and their interactions with the microenvironment [13]. Different SRT protocols exist, with trade-offs between cellular and transcriptomic resolution [14]. For example, MERFISH detects transcripts in single cells with high efficiency but only for a targeted panel of genes [15], while Visium HD measures the whole transcriptome very sparsely in each cell [16, 17]. In contrast, the widely used Visium platform measures the whole transcriptome in spots of 55 *µ*m in diameter that aggregate transcripts from up to 10 cells [18], which makes high-resolution cell states and cancer clones difficult to identify from these data alone.

To address these limitations, researchers have developed computational methods to integrate single-cell RNA-sequencing (scRNA-seq) with spot-based SRT data from the same tissue. These methods either deconvolve the gene expression of each spot into reference cell types identified in scRNA-seq [19, 20], map individual cells to spots [21], or both [22].

The resulting integration enables the mapping of highly-resolved cell states in the TME [23] and the identification of spatial niches [12, 24]. The underlying principle among these methods is that the cell populations of interest have distinct gene expression profiles, which enable their identification in both modalities. However, these differences are often not apparent among subclones of a tumor, as their phenotypes are only partly heritable and change due to cell state plasticity [4, 25]. Therefore, the same assumptions that enable cell type mapping in SRT through integration with dissociated single-cell data do not permit the accurate identification of cancer clones, preventing an accurate study of spatial tumor evolution.

In contrast, cancer clones may be identified in transcriptomics data from somatic mutations. In scRNA-seq, this may be achieved through copy number aberration (CNA) calling directly [26], or by integration with DNA sequencing data [27, 28], which may even enable phylogenetic tree inference from single-nucleotide variants (SNVs) [29]. Similar approaches have been developed for SRT data, including CNA calling [30–32] and mapping SNVs identified in bulk DNA-seq to spatial spots [33]. In methods for SRT, the spatial context can partly compensate for the lower data resolution, ascells from the same subclone will tend to have similar spatial locations. However, the subclones and phylogenies inferred from SRT data remain coarser than those obtained from dissociated single cells, as the signal of each subclone is diluted by the other cells in each spot [30], and in CNA-based methods each spot is assigned to a single subclone [30, 32]. Unlike methods to map cell types in SRT data, methods to identify subclones have not yet leveraged the higher resolution subclonal signal that scRNA-seq data may provide.

Here, we present SpOTr, an optimal transport (OT) [34] and spot deconvolution approach to map both the cell types of the TME and the cancer clones from dissociated scRNA-seq to spot-based SRT data, using the phylogenetic tree that relates the cancer clones. SpOTr is based on the assumption that evolutionarily related cancer cells tend to be in spatial proximity within a tumor, and the tree therefore contains information about the locations of the subclones that cannot be determined solely by their gene expression profiles. SpOTr is the first method to integrate scRNA-seq and SRT data that combines spot deconvolution based on cell state signatures, with Fused Gromov–Wasserstein (FGW) OT [35] based on the concordance of an evolutionary tree of cancer cells and their spatial locations. The method iterates between optimizing the parameters of the spot deconvolution model for both TME cell types and cancer clones, and computing the mapping of individual cancer cells to spots. This enables the identification of cancer clones in spatial data without requiring CNA or SNV calling from spots, and it accounts for the presence of different TME cell type compositions in spots, which are mapped simultaneously.

In the following sections, we first provide an overview of SpOTr. We then benchmark SpOTr on simulated tumors, and obtain more accurate clone compositions of the spots than spot deconvolution and cell mapping methods. Next, we apply SpOTr to synthetic Visium spots constructed from PEtracer lineage-traced tumors [9], and place the clades in better agreement with their in situ readout than competing methods. Finally, we apply SpOTr to a human colorectal liver metastasis [36], and find that using the concordance between tree and spatial structure enabled the identification of two subclones that were not obtained by expression-only deconvolution.

## Results

### Method overview

SpOTr integrates scRNA-seq data, a phylogenetic tree of the cancer cells, and spot-based SRT data from a section of the same tissue (Figure 1). To account for the mixture of cells in each spot, SpOTr uses the spot deconvolution model of cell2location [20]. In this model, the counts of each gene in each spot follow a negative binomial distribution with mean defined by the contribution of different cell type signatures weighted by their abundances in the spot. SpOTr separates these contributions into the TME cell types and the cancer clones, where the abundances of the clones are obtained by a mapping *γ* of the cancer cells to the spots.

**Figure 1.**
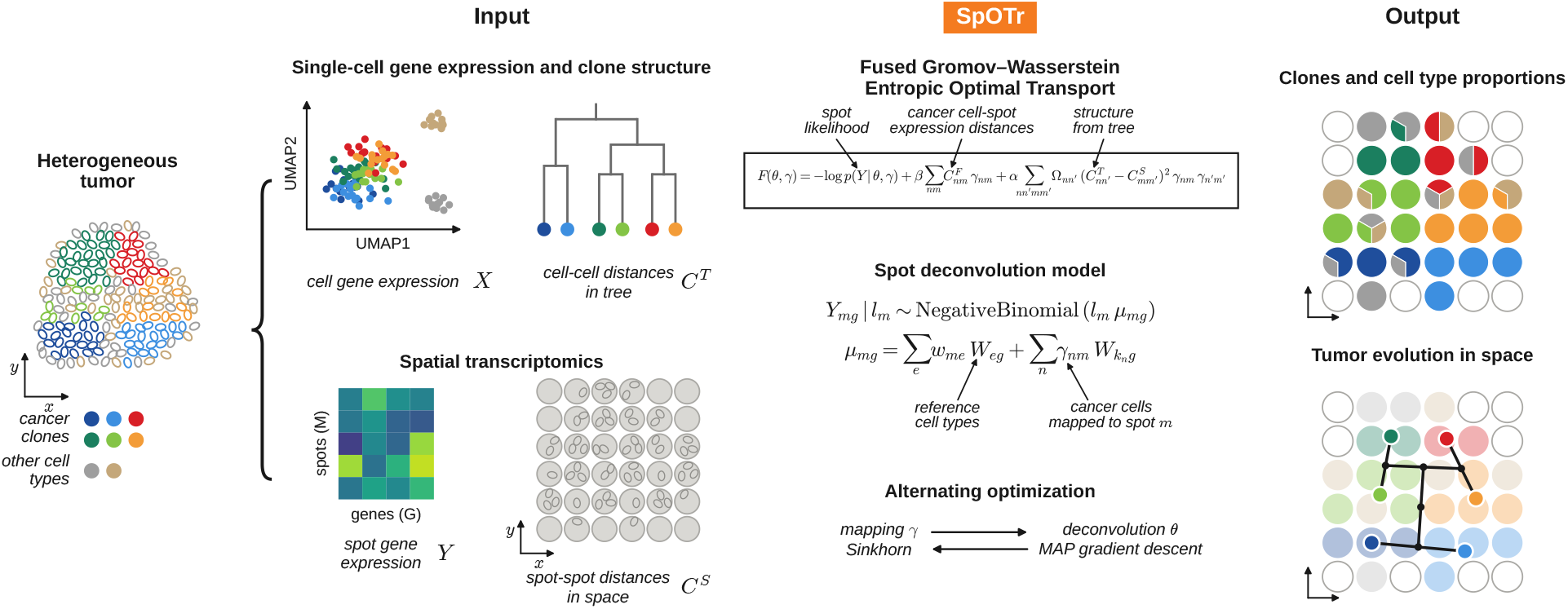
SpOTr overview. SpOTr takes as input scRNA-seq data of dissociated cells of every cell type, with a phylogenetic tree relating the cancer cells, and spot-based SRT data with the positions of the spots. The cancer cells are mapped to the spots by entropic optimal transport with an FGW cost that combines the likelihood of the spots, the distances between the expression of cells and spots, and a structural term. The structural term matches the cell-cell distances in the tree *C*^*T*^ to the spot-spot distances *C*^*S*^, with optional weights for each clade in Ω. The optimization alternates between the mapping *γ*, updated by the Sinkhorn algorithm, and the parameters *θ* of the deconvolution model, updated by gradient descent to their maximum a posteriori (MAP) estimate. The likelihood of the spot gene expression combines cell types with the cancer cells mapped to it, each with the signature 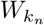 of its clone *k*_*n*_. The outputs are the cell type and clone composition of each spot and the location of each cancer cell, which can be used to study spatial tumor evolution.

SpOTr formulates the spatial mapping of subclones as an OT problem with an FGW cost that combines three terms (Methods). The first is the negative log-likelihood of the spots under the deconvolution model adapted from cell2location, the second is the cosine distance between the normalized expression profiles of each cancer cell and of each spot, and the third is a structural term that compares the evolutionary relationships between cells with the spatial distances between the spots they are mapped to. In this OT formulation, the number of cancer cells that map to each spot is informed by the deconvolution, and therefore by the presence of TME cell types.

Crucially, the structural term matches the distances between cells in the tree to the spatial distances between spots, which draws closely related cells to nearby spots and distantly related ones apart. This term can be seen as a regularization of the expression-only OT cost, allowing cells from subclones with poor gene expression specificity to be assigned to a spatially coherent neighborhood. When only clone labels are available, the term instead draws the cells of each clone together without constraining how far apart the clones lie (Methods). Finally, SpOTr also supports the incorporation of prior knowledge about the spatial organization of specific clones (Methods, Figure S5).

The optimization alternates between computing the mapping with the Sinkhorn algorithm [37] on a linearization of the objective (Supplementary Note 1) and taking gradient steps on the parameters of the deconvolution model (Methods). SpOTr then outputs the cell type and clone composition of each spot and the location of each dissociated cancer cell. These can be used to study how the tumor evolved in space, for example by drawing the tree in the tissue.

### Benchmarking study on simulated data

We assess the ability of SpOTr to recover the clone composition of each spot in the setting it is designed for, in which the clones are hard to distinguish by their gene expression but each occupies its own region of the tissue (Figure 2A, B). Specifically, we simulate the lineage of each tumor with the birth-death process of Cassiopeia [38] and record it with Cas9 lineage tracing. We place the cells in the tumor nests of a microenvironment simulated by scCube [39], with each clade of the tree in a contiguous territory within that of its parent. We simulate gene expression from heritable programs that evolve along the tree, weak positional programs that vary across the tissue, and programs for three synthetic non-tumor cell types. We then measure one half of each tumor with Visium-like spots, which hold about 12 cells each and miss the cells between them (Figure S2). From the other half, we sample dissociated cells, and the two modalities thus share clones but not cells (Methods). Starting from this setting, we vary how much the clones differ in their expression, how many of them are dispersed over the section, and how many clones there are.

**Figure 2.**
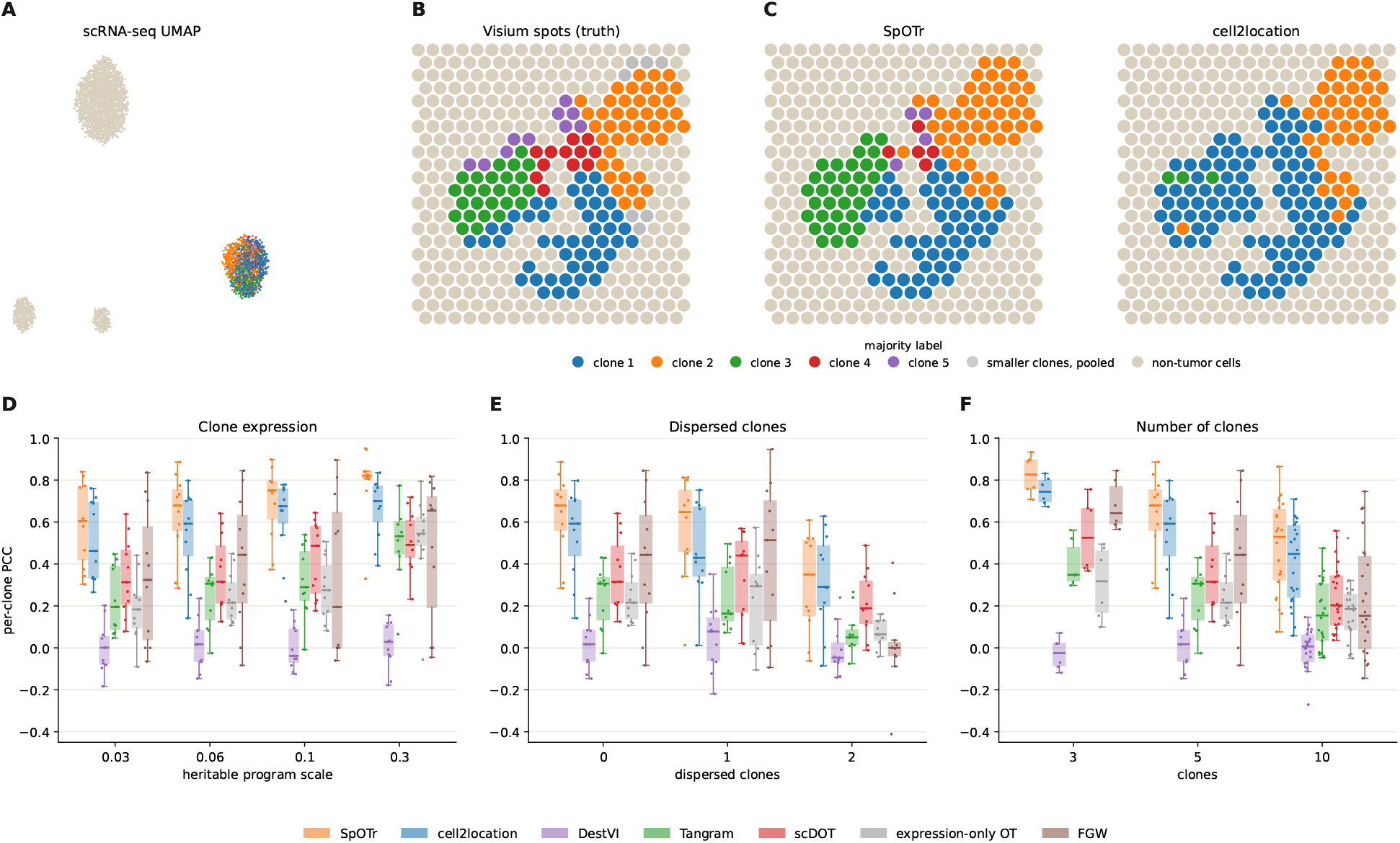
Benchmarking study on simulated data. (A) UMAP of the gene expression of the dissociated cells of a simulated tumor, with the tumor cells colored by clone and the non-tumor cells in beige. The clones are hard to distinguish by their expression. (B) The Visium spots of the other half, each colored as non-tumor where non-tumor cells outnumber tumor cells, and otherwise by the majority clone of its tumor cells. Each clone holds its own territory. The lineage comes from Cassiopeia and the tumor nests and microenvironment from scCube. (C) The same for SpOTr and for cell2location, the most accurate of the other methods. (D–F) The per-clone Pearson correlation between the true and the estimated clone proportions of the spots, as the strength of the heritable programs (D), the number of clones whose cells are dispersed over the section (E) and the number of clones (F) vary. The others are as in A and B, and the clones of both seeds are pooled. SpOTr is run with the lineage tree. Figure S1 shows the other metrics.

We run SpOTr with the lineage tree and compare it with cell2location [20], DestVI [19], Tangram [21] and scDOT [22]. To separate the contributions of the structural term and of the deconvolution model, we also include exact optimal transport on gene expression alone and FGW optimal transport [35] with the same distance matrices as SpOTr. We score each method by the Pearson correlation between the true and the estimated proportion of each clone across the spots (Figure 2D–F), and report the structural similarity, the root mean squared error and the Jensen–Shannon divergence [40] in Figure S1.

On an example tumor, SpOTr recovered the territories of the three largest clones and part of the two smallest, while cell2location assigned two thirds of the tumor spots to the largest clone and almost none to the three smaller ones (Figure 2B, C). Over all simulated tumors, SpOTr recovered the clone composition of the spots better than all other methods in every scenario (Figure 2D–F), and on each of the four metrics (Figure S1). cell2location, which deconvolves the spots with each clone as a reference cell type, was the most accurate of the other methods (Figure 2D–F). Tangram and exact optimal transport, which map cells by their expression alone, placed the clones poorly, since the clones cannot be told apart from the expression of single cells in this setting. In contrast, the advantage of SpOTr was largest when the clones were hardest to distinguish, and it remained the most accurate when they differed clearly in their expression (Figure 2D).

FGW uses the same tree and expression as SpOTr, but it improved less over expression alone. Without a deconvolution model, FGW gives every spot the same capacity, and it placed much of the mass of the cancer cells in spots without tumor cells. In contrast, SpOTr placed almost all of it in tumor spots. Even within the tumor spots, SpOTr was more accurate than FGW in every simulated tumor, since the likelihood of the spots depends on how well the clone signatures explain the cancer part of each spot. scDOT placed the clones better than expression alone but less accurately than SpOTr. DestVI recovered the tumor fraction of each spot but did not separate the clones in any scenario.

As expected, the advantage of SpOTr shrank when clones were dispersed over the section, where the structural term draws together cells that lie apart (Figure 2E). With two of five clones dispersed, its median correlation fell to 0.35, close to that of cell2location (0.29; Figure 2E). When the dispersed clones are known, setting their weights in the structural term to zero improved the per-clone correlation of the clones dispersed at random in nine of ten cases, by up to 0.13, without changing the other clones (Figure S5). It lowered the correlation of the one dispersed clone that remained clustered in the tissue, for which the structural term was still informative. With more clones, all methods lost accuracy, and SpOTr kept its lead over cell2location, with median correlations from 0.83 against 0.74 with three clones to 0.53 against 0.45 with ten (Figure 2F).

### Application to PEtracer lineage-traced tumors

PEtracer [9] records the lineage of each cell with an evolving barcode that is read both in dissociated cells and in situ. This provides a ground truth for the location of each clone in a tumor section, and makes a semi-synthetic benchmark with real cells at their real positions and synthetic Visium spots built from them.

The cells of section M3-4 with a lineage readout form clades that occupy distinct regions of the tumor, and the synthetic Visium spots keep this arrangement (Figure 3A, B). Since only 31% of the tumor cells carry a readout (Figure 3A), we scored each method on the proportions of the clades in each spot, which are known for these cells. SpOTr recovered the main regions of the clades, keeping the clades of the upper part of the tumor apart from those of its lower part as in the truth (Figure 3C). In contrast, cell2location correctly placed the clades of the upper part but assigned much of the lower part to a few clades, and DestVI assigned most tumor spots to a single clade (Figure 3D, E). Tangram and scDOT spread the clades over the whole tumor (Figure 3F, G). As in our simulations, the PEtracer clades are poorly separated by their expression, since the nearest expression neighbors of each cell often belong to other clades. Accordingly, SpOTr agreed best with the truth on all four metrics, clade by clade (Figure 3H–K).

**Figure 3.**
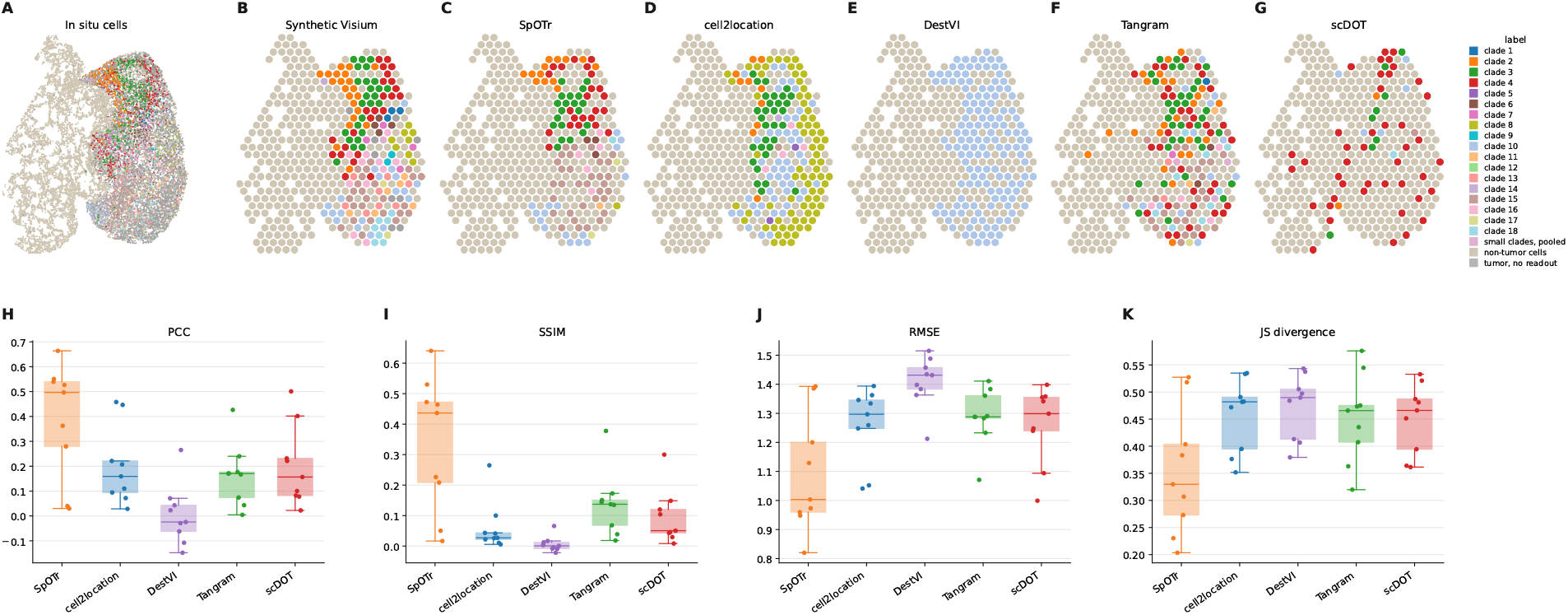
Application to PEtracer lineage-traced tumors. (A) The cells of one half of section M3-4 at their in situ positions, each tumor cell colored by its clade from the in situ lineage readout, the tumor cells without a readout in gray and the non-tumor cells pooled. (B) The synthetic Visium spots built from these cells. Each spot is colored as non-tumor if it has more non-tumor than tumor cells, and otherwise by the majority clade among its tumor cells with a lineage readout, which 31% of tumor cells have. (C–G) The same for the maps of SpOTr with the lineage tree, cell2location, DestVI, Tangram and scDOT. (H–K) Per-clade agreement with the truth, the Pearson correlation and the structural similarity (higher is better), and the root mean squared error and the Jensen–Shannon divergence (lower is better) between the true and the estimated clade proportions of the spots, with one point per clade.

### Application to a human colorectal liver metastasis

The Human Tumor Atlas Network (HTAN) case HT260C1 [36] is a colorectal liver metastasis with single-nucleus multiome data and a Visium section from the same tissue block. We identified two main clones, A and B, and two sub-lineages of clone B, B1 and B2, from the CNA signals of the nuclei in their chromatin accessibility and in their gene expression, which we computed as in inferCNV [41] (Methods). Here we focus on cell2location as a baseline, which uses the clones as reference cell types and estimates how many cells of each clone each spot holds, but it cannot place individual cells or resolve structure within the clones it is given.

The CNAs of the nuclei separate clones A and B and the sub-lineages B1 and B2, and the tree built from them groups the nuclei of each clone (Figure 4A). SpOTr and cell2location found the same overall arrangement of the tumor, with clone A on the right side of the section and clone B in the nests on its left (Figure 4B, C). However, SpOTr separated B1 from B2 more clearly than cell2location, as measured against the CNA signal of held-out genes (Figure S3). In addition to the clones it is given, SpOTr placed the two sub-lineages of B1 in agreement with their CNA signal in the Visium data, with B1b in the nests on the left and B1a in the upper band of the section (Figure 4D, E). By placing each lineage at the centroid of its cells, we are able to draw the tree in the tumor slide, from the common ancestor near clone A to the nests of clone B and the separate regions of B1a and B1b (Figure 4F).

**Figure 4.**
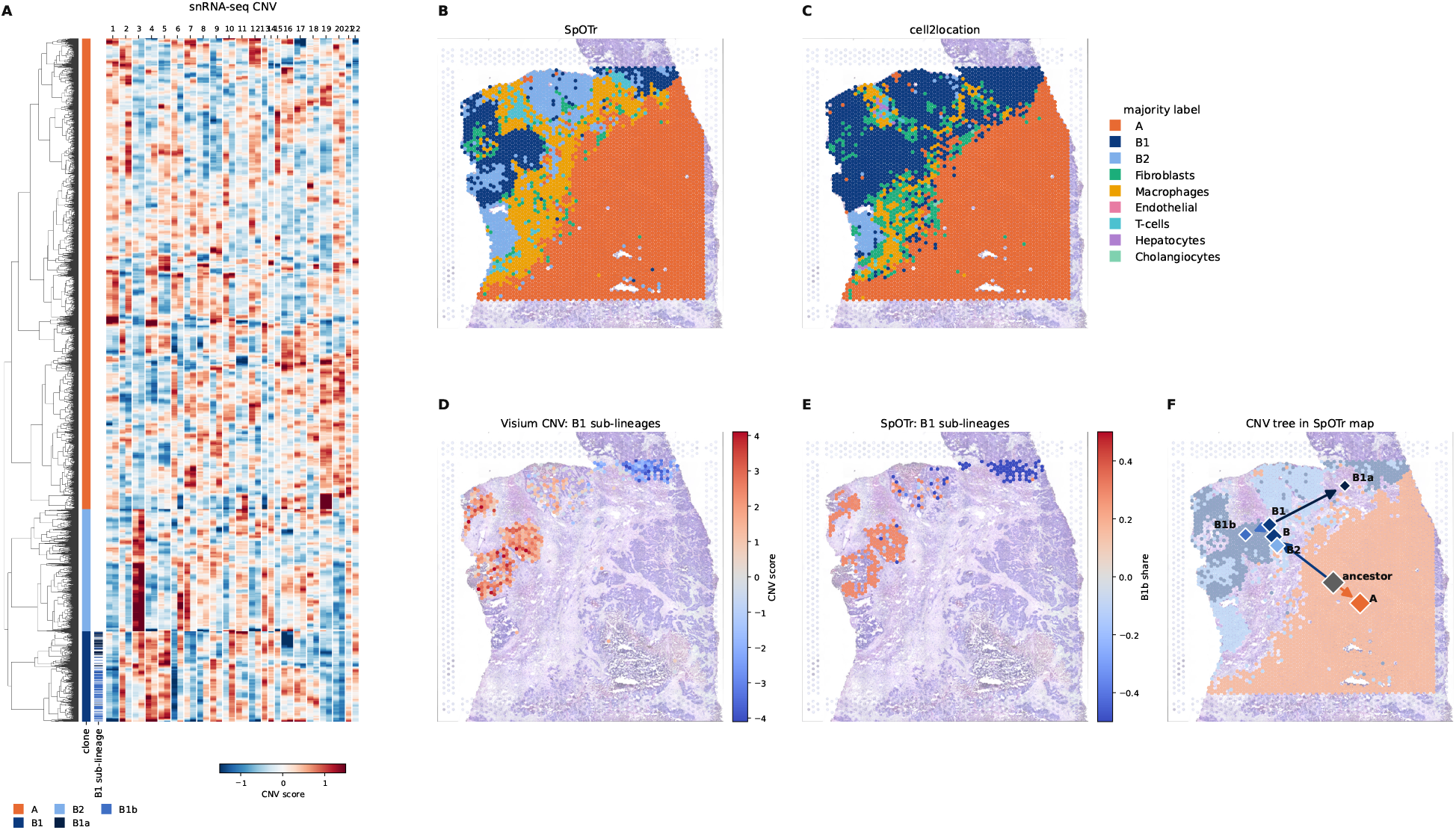
Application to a human colorectal liver metastasis (HT260C1 of the Human Tumor Atlas Network [36]). (A) The CNA profiles of the cancer nuclei from their gene expression, computed as in inferCNV, ordered by the tree that SpOTr uses, with the clone of each nucleus and the two sub-lineages of clone B1. (B, C) Each spot colored by its majority label across all cell types and clones, for SpOTr with the tree and for cell2location. (D) The CNA signal of the two sub-lineages of B1 in the Visium data, which no method is given as labels, inside the territory of B1 (B1b in red), and (E) the share of B1b among the cells of B1 that SpOTr maps to each spot, relative to the share of B1b among all nuclei of B1. (F) The tree in space, with each clone, sub-lineage and ancestor at the centroid of its nuclei under the mapping of SpOTr, and arrows from each ancestor to its descendants. The agreement of each mapping with the CNA signal is shown in Figure S3.

## Discussion

The locations and functional states of the cells within a tumor shape its evolutionary dynamics. Resolving this heterogeneity requires the integration of data modalities that together identify not only the different cell types and their spatial locations, but also the locations of cancer clones and their gene expression profiles. Here we developed SpOTr, the first method to couple spot deconvolution with an OT mapping informed by the phylogeny of the cancer cells, which maps both the cancer clones and the TME cell types from scRNA-seq data to spots while promoting the spatial coherence of related cancer cells.

Our results show that SpOTr maps cancer cells better than methods that rely on gene expression alone. In simulations, SpOTr recovered the clone composition of the spots more accurately than competing methods, with the largest advantage when the clones were transcriptionally similar. On a PEtracer section [9], whose in situ lineage readout gives the true location of each clade, SpOTr recovered the clade composition of its synthetic spots. In HT260C1 [36], which has no lineage readout but represents a realistic application scenario for our method, SpOTr placed subclones that no method is given as labels in agreement with the CNA signal of the spots.

SpOTr can be seen as an extension of cell2location [20] in which the mapping of the cancer cells is guided by the spatial coherence of their clones. Like scDOT [22], it couples spot deconvolution with a cell-level mapping, but scDOT works in two steps, deconvolving the spots by non-negative least squares and then fitting an OT mapping of the cells to these proportions, whereas SpOTr learns the mapping and the deconvolution jointly. TemSOMap [42] also uses the lineage of dissociated cells to map them to spatial data, but it assumes that every dissociated cell is part of the lineage tree, and it cannot map the cancer clones and the TME cells together. Its code was not publicly available, and we could not include it in our comparisons. Finally, in contrast to methods that call CNAs from SRT spots [30, 32], SpOTr takes the clones and their phylogeny from the dissociated cells, where they can be resolved more finely, and requires no CNA or SNV calling in the spots.

SpOTr assumes that each clone is represented in the dissociated sample in the same proportion as in the section, which may not hold when the two samples come from different regions of a tumor. Learning the total of each clone in the section from the spots, with the composition of the dissociated sample as a prior, would relax this assumption. The structural term also assumes that related cells lie close together, and its importance *α* is set by the user rather than learned from the data. The elbow of the trade-off between the expression and structural costs can guide this choice (Figure S4), but it does not indicate whether the tree carries spatial information, and when clones disperse over the section, a high *α* pulls their cells together incorrectly. Setting the weights of such clones to zero improved their maps in our simulations (Figure S5), and learning these weights from the data, as well as propagating the uncertainty of the tree [38], are natural extensions of SpOTr.

SpOTr is not limited to cancer subclones related by a phylogenetic tree. Clones that are known without their relationships, such as T and B cell clones grouped by their receptor sequences or clones marked by static barcodes such as LARRY [43], CellTag [44] and TREX [45], can enter as labels, which draws the cells of each clone together. In principle, spatial transcriptomics at cellular or sub-cellular resolution does not require spot deconvolution. In practice, however, sequencing-based platforms such as Visium HD capture so few transcripts in each 2 *µ*m bin that the bins are merged into bins of 8 *µ*m by default, which do not match the cells and are themselves deconvolved [16, 46], and SpOTr applies to such bins as it does to spots. Imaging-based platforms such as MERFISH do resolve single cells, but only for a panel of genes [15], and there the mapping of SpOTr could be used without the deconvolution model.

Data integration is an important step in cancer studies, but the complexity of the underlying biology and the unique features of different data modalities make this task a major computational challenge. By leveraging the unique benefits of each modality, and incorporating prior knowledge about their relationship explicitly, SpOTr and its future extensions will enable researchers to characterize the spatial organization of tumor clones and the TME from their own data sets.

## Methods

### Notation

Let the SRT data consist of the UMI counts of *G* genes in *M* spots, *Y* ∈ ℕ^*M ×G*^, with *l*_*m*_ ∈ ℝ^2^ the position of spot *m* and 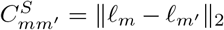 the distance between spots *m* and *m*^*′*^. For the scRNA-seq data, we consider TME cells with cell types *e* ∈ *{*1, … , *E }*, and *N* cancer cells with UMI counts *X* ∈ ℕ^*N ×G*^ that belong to clones *k*_*n*_ ∈ *{*1, … , *K}* . The cancer cells are related through a phylogenetic tree, and 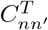 is the distance between cells *n* and *n*^*′*^ in the tree. Bold symbols denote vectors, such as the counts **y**_*m*_ of spot *m* and **x**_*n*_ of cancer cell *n* over the genes.

### Spot deconvolution model

SpOTr models the counts of each spot with the negative binomial model of cell2location [20], with an additional term for the cancer cells mapped to the spot,

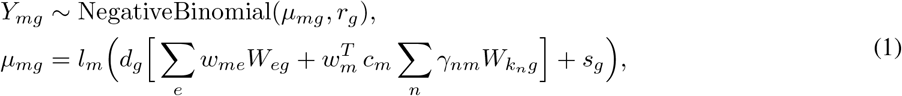

where *W*_*eg*_ and 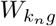 are the reference signatures of cell type *e* and of clone *k*_*n*_, *w*_*me*_ and 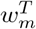 are the numbers of cells of type *e* and of cancer cells with a clone in spot *m, γ*_*nm*_ is the mass of cancer cell *n* that the mapping transports to spot *m, d*_*g*_ is the sensitivity of gene *g, l*_*m*_ is the detection efficiency of spot *m, s*_*g*_ is an additive background and *r*_*g*_ is the dispersion of gene *g*. The expected counts of spot *m* are thus the sum of the counts expected from its microenvironment cells and from its 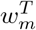 cancer cells, and the cancer cells express 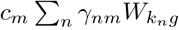 , the mean clone signature of the cells mapped to the spot, weighted by their masses *γ*_*nm*_, and normalized by the total number of cancer cells mapped to spot *m*, with *c*_*m*_ = 1*/∑* _*n*_ *γ*_*nm*_.

The contributions of the detection efficiency *l*_*m*_, the gene sensitivity *d*_*g*_ and the additive background *s*_*g*_ to the counts follow the cell2location model. The signatures are estimated from the dissociated cells with the regression model of the cell2location package, one for each cell type of the microenvironment and one for each clone. However, while cell2location would give each clone its own abundance in each spot, the abundance of clone *k* in spot *m* is here 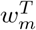 times the fraction of the mass mapped to the spot that comes from clone *k*. Cancer cells without a clone, such as tumor cells without a lineage readout in lineage tracing experiments, enter as one more reference type with its own signature and abundance, as in cell2location. For simplicity, in the following we assume that every cancer cell is in the phylogenetic tree.

Each spot holds *N*_*m*_ cells, and their composition, the fractions of cells of each type and of cancer cells, is the softmax of unconstrained parameters **z**_*m*_ ∈ ℝ^*E*+1^. The numbers of cells and the other parameters of Equation (1) follow

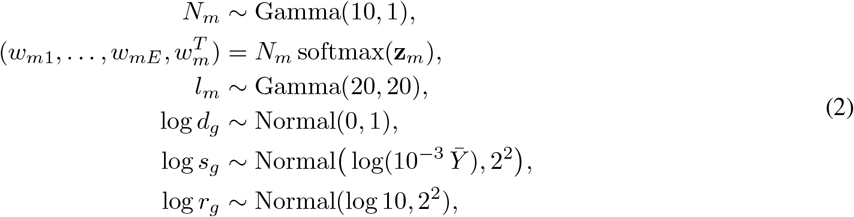

where 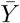 is the mean count over all spots and genes. The priors on *N*_*m*_ and *l*_*m*_ are those of cell2location. The prior on *N*_*m*_ is its prior on the number of cells per spot, with the mean set to the 10 cells that a Visium spot can hold [18], and the prior on *l*_*m*_ is its default prior on the detection efficiency, with mean 1. The other priors are weakly informative, with standard deviations of 1, 2 and 2 on the log scale. They center the sensitivity of each gene at 1, as in cell2location, the background at a small fraction of the mean count, and the dispersion at 10.

### The mapping and its capacities

The marginals of the OT mapping, the total mass that it moves from each cancer cell and into each spot, tie the deconvolution and the mapping together, and enable the deconvolution to set how many of the dissociated cancer cells each spot receives. The mapping 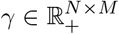 is a transport plan in which *γ*_*nm*_ is the mass of cancer cell *n* placed in spot *m*, with row marginals *a*_*n*_ = ∑_*m*_ *γ*_*nm*_ and column marginals *b*_*m*_ = ∑_*n*_ *γ*_*nm*_. Each cancer cell carries mass 1, *a*_*n*_ = 1, and the total mass is therefore *N* . The column marginal *b*_*m*_ is then the number of dissociated cancer cells mapped to spot *m*, and we call it the capacity of the spot. The factor *c*_*m*_ in Equation (1) is its inverse, *c*_*m*_ = 1*/b*_*m*_. The capacities divide the *N* cancer cells among the spots in proportion to the share of the cells of each spot that the deconvolution (Equation (1)) counts as cancer cells,

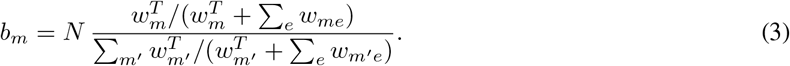

The capacities follow the share of cancer cells rather than their number *w*^*T*^ , because the number of cells in a spot trades off with its detection efficiency *l*_*m*_ and is only weakly determined by its counts, whereas its composition is not. Because of the column marginals, cancer cells are only mapped to spots that the deconvolution assigns cancer cells to, unlike in FGW with uniform marginals. Because of the row marginals, each clone keeps its share of the dissociated cells over the whole section, which assumes that the dissociated sample represents the clones of the section in proportion.

### Constructing the SpOTr objective

SpOTr combines the deconvolution model in Equation (1) with FGW transport to estimate the mapping *γ*. The FGW terms compare the expression of cells and spots and relate the phylogeny of the cancer cells to their spatial organization.

#### The expression cost

The expression cost 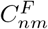 compares the expression of cancer cell *n* with the expression of the cancer cells of spot *m*, and is the feature cost of FGW [35]. The expression of the cancer cells of spot *m* is estimated by its counts less those that the deconvolution assigns to its microenvironment cells and to the background,

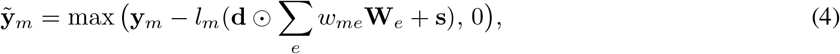

where ⊙ denotes the elemeLntwise product. With the log-normalized expression of cell *n* and of the cancer cells of spot *m*, **u**_*n*_ = log(1 + 10^4^ **x**_*ng*_ */∑*_*g*_ *x*_*ng*_) and 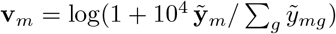 , the expression cost is their cosine distance,

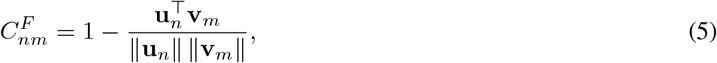

which is also the starting cost of scDOT [22] and the similarity of Tangram [21], and does not depend on sequencing depth. Because **v**_*m*_ depends on the deconvolution, the cost is recomputed after each update of *θ*.

#### The structural term

The structural term compares the relationships between cancer cells with the distances between the spots they are mapped to,

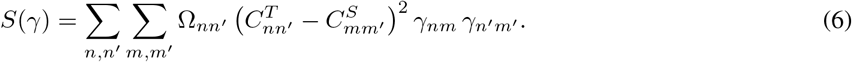

Each pair of cancer cells *n* and *n*^*′*^ placed in spots *m* and *m*^*′*^ contributes the squared difference between their distance 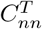 and the distance 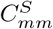 between their spots, weighted by 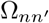 and by the masses *γ*_*nm*_ and 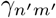 . The term is small when the distances between the spots of the cells agree with the distances between the cells. The pair weights 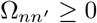 , normalized to sum to *N* , set which pairs of cells must show this agreement and how strongly. A pair with 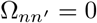 may lie anywhere relative to each other, and the larger 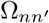 , the more closely the distance between the spots of the two cells must match their distance 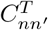 .

Together, Ω and *C*^*T*^ encode what is known about how the cancer cells are related. In practice, this is often either just the clone of each cell or a phylogenetic tree of the cells. Clone labels alone are available when clones are known without their relationships, as for clones marked by static barcodes, and a tree when the lineage is recorded by evolving barcodes or reconstructed from SNVs or CNAs. We first describe the structural term with clone labels, the simpler case, and then what a tree adds.

When only the clone of each cell is known, 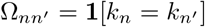 and *C*^*T*^ = 0. Up to the constant that normalizes Ω, Equation (6) is then the sum over clones of the squared distances between the spots of their cells,

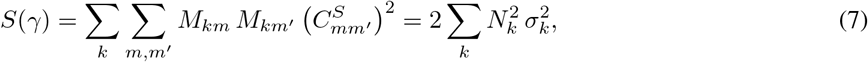

where 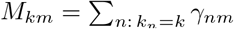 is the mass of clone *k* in spot *m, N*_*k*_ = ∑_*m*_ *M*_*km*_ is the number of cells of clone *k*, and 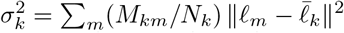 is the spatial variance of clone *k* around its center 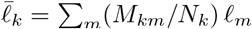 . This scenario therefore draws the cells of each clone together, and since pairs of cells from different clones have zero weight, it does not constrain how far apart different clones lie.

When the cells are related by a phylogenetic tree, 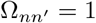 for every pair, and 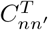 is the length of the path between cells *n* and *n*^*′*^ in the tree, rescaled to have the same mean over pairs as the distances between spots. Pairs of cells from different clones then also enter the term, and closely related cells are drawn together and distantly related ones apart. The tree thus also places the clones relative to each other, which clone labels leave unconstrained, and sister lineages of one clone may occupy separate regions.

Users can weight the term for each clade, 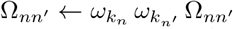 , where *ω*_*k*_ = 0 maps clade *k* by expression alone and *ω*_*k*_ *<* 1 relaxes it, for example for a clade known to infiltrate the tissue (Figure S5). By default, *ω*_*k*_ = 1.

Linearized at the current mapping, the structural term gives the cost of placing cell *n* in spot *m* as the masked Gromov–Wasserstein cost derived in Supplementary Note 1,

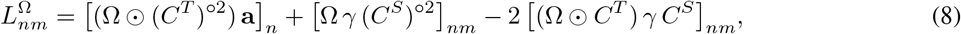

where **a** = *γ***1** is the vector of row sums of the mapping and *A*°^2^ the elementwise square of a matrix *A*. With 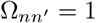 for every pair, Equation (8) is the linearized cost of Gromov–Wasserstein [47].

#### The SpOTr objective

Let *p*(*Y* | *θ, γ*) and *p*(*θ*) be the likelihood and prior functions of the deconvolution model in Equation (1) and eq. (2). SpOTr jointly estimates the parameters *θ* of the deconvolution model, *N*_*m*_, **z**_*m*_ and *l*_*m*_ for every spot and *d*_*g*_, *s*_*g*_ and *r*_*g*_ for every gene, and the mapping *γ*, by minimizing

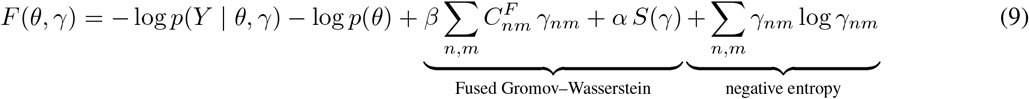

over *θ* and over the mappings with the marginals above, where *β* ∈ [0, 1) and *α*∈ [0, 1) are the importances of the expression cost and of the structural term relative to the likelihood. We weigh the two FGW terms against the likelihood rather than against each other as in the usual FGW [35], so that *α* changes only the structural term. For a fixed *θ*, this is an entropic FGW problem [35] with an additional negative log-likelihood regularizing term. The cost involves two linear-complexity terms (the negative log-likelihood of the spots and the expression cost *C*^*F*^), while the structural term has quadratic complexity.

#### Strengths of the terms

The likelihood, the expression cost and the structural term are measured in different units. At the start of the optimization, we take the gradient of each term at the initial mapping, a cost per cell and spot, and measure its spread by the standard deviation of this cost after subtracting its row and column means, which do not change the transport plan. We then scale the expression cost and the structural term to *β/*(1 *− β*) and *α/*(1 *− α*) times the spread of the negative log-likelihood, and *β* and *α* in Equation (9) stand for these calibrated multipliers. We use *β* = 1*/*2 and *α* = 0.2 throughout, so the two costs keep the usual FGW proportion of 1 *− α* to *α*. The per-clone weights apply after this calibration, which leaves the strength of the other clones unchanged. On new data, the elbow of the trade-off between the expression cost and the structural cost over a grid of *α* can guide its choice (Figure S4).

### Inference

SpOTr alternates between the mapping and the deconvolution, updating each given the current estimate of the other. To update *γ* given *θ*, the objective is linearized into one cost per cell and spot, consisting of the gradient of the negative log-likelihood plus the weighted expression cost plus the weighted linearized structural cost of Equation (8). The entropic optimal transport plan of this cost under the marginals above is computed by Sinkhorn iterations in the log domain [37]. The mapping then moves towards this plan by a step chosen on a grid from 1 to 0.001, each step scored by its exact change of the objective. Given *γ*, the parameters *θ* are updated by Adam [48] on the log-likelihood and the priors, effectively pursuing a maximum a posteriori estimate of *θ*, with the expression of the cancer cells of each spot fixed to the mean signature of the cells mapped to it. The mapping thus shapes the deconvolution through the clone composition of each spot, which changes how much expression the microenvironment must account for and how many cancer cells each spot holds in the next update. The model is written in JAX [49], and the schedule of the updates is given in the Supplementary Methods.

### Alternative methods

Cell2location [20] and DestVI [19] deconvolve each spot into reference cell types, with every clone and every other annotated cell type as a reference type. Tangram [21] maps the dissociated cells to the spots by the similarity of their gene expression, and scDOT [22] couples a deconvolution of the spots with an optimal transport map of the cells, from which we read out the map. On simulated data, we also compare with exact optimal transport between all dissociated cells and the spots on the expression cost of SpOTr, and with Fused Gromov–Wasserstein optimal transport [35] with the same distance matrices as SpOTr and uniform marginals. Tangram and exact optimal transport are given the true number of cells in each spot, which SpOTr estimates instead.

### Simulations

Each scenario varies one property of the tumor, the strength of the heritable programs (0.03, 0.06, 0.1 and 0.3), the number of clones whose cells are dispersed over the section (none, one or two) or the number of clones (three, five or ten), with the others at their middle level, and each setting is simulated with two seeds. We chose the settings of SpOTr, *α* = 0.2 and *β* = 1*/*2, on a third seed of the middle setting, which the benchmark does not score, and used them throughout. The Supplementary Methods describe the simulation in full.

### Real data

For the PEtracer data, the time-calibrated tree, clades and cells are those of [9]. In each section, synthetic Visium spots are built from a random half of the cells, and 8,000 cells drawn from the other half form the dissociated sample. For the HTAN data, we use case HT260C1 of [36], a colorectal liver metastasis with single-nucleus multiome data and a Visium section from the same block. All methods fit one random half of the genes, and the CNA signal of the spots, which serves as a reference, is computed from the other half.

## Code availability

SpOTr is freely available as a Python package in https://github.com/pedrofale/spotr. The code repository also contains a Snakemake [50] workflow and Python scripts to reproduce all the results shown in this paper.

## Data availability

The PEtracer data [9] were obtained with pycea 0.2.0 (https://github.com/YosefLab/pycea) from https://zenodo.org/records/15750529. The colorectal liver metastasis data [36] were downloaded from the Human Tumor Atlas Network data portal (https://data.humantumoratlas.org).

