## Supplementary Information for "Phylospatial optimal transport for single-cell and spatial transcriptomics data integration"

[1] *Division of Computational Science and Technology, KTH, Royal Institute of Technology*
[2] *SciLifeLab*
[3] *Department of Cell and Molecular Biology, Karolinska Institutet*
[4] *Department of Biosystems Science and Engineering, ETH Zürich*
[5] *Department of Computer Science, ETH Zürich*
[6] *ETH AI Center*

#### Supplementary Methods

##### Method settings

Cell2location [1] and DestVI [2] deconvolve each spot into reference cell types, and we use every clone and every other annotated cell type as a reference type. We train the reference model of cell2location for 250 epochs and its spatial model for 30,000 epochs with a detection alpha of 20. We set its prior on the number of cells per spot to the mean true number of cells in each spot in the simulations and in PEtracer and to 10 on HT260C1. We train the single-cell model of DestVI for 300 epochs and its spatial model for 1,500 epochs. Tangram [3] maps the dissociated cells to the spots by the similarity of their gene expression, and we run it in its cell mode for 1,000 epochs with the true number of cells in each spot as its density prior. scDOT [4] couples a deconvolution of the spots with an optimal transport map of the cells, and we read out its map after the ten iterations of its documented training loop. We also compare with two baselines from POT [5] on simulated data. Exact optimal transport maps all dissociated cells to the spots on the cosine distance between log-normalized expression profiles and uses the true number of cells in each spot as the mass of the spot. Fused Gromov–Wasserstein optimal transport [6] maps the lineage-traced cells on the same cosine distance and on the tree distances given to SpOTr and the distances between the spots. We divide the last two by their maxima and use uniform marginals and equal weights on the expression and the structure. Both baselines use the whole expression of each spot, whereas the expression cost of SpOTr uses the expression of the cancer cells of the spot (Methods). We run SpOTr with the lineage tree in JAX [7]. It first fits the parameters of the deconvolution for 1,000 steps of Adam with every cancer cell spread over the spots in proportion to their capacities, and then alternates 16 times between 12 updates of the mapping and 200 steps of Adam.

##### Simulations

We simulate each tumor with Cassiopeia [8]. We simulate the lineage of about 12,000 tumor cells as a birth-death process with weak selection, in which fitness changes at rate 0.2 with standard deviation 0.1. We record the lineage by Cas9 lineage tracing at 13 cassettes of three target sites. We take as clones the five clades that contain at least 3% of the cells at the earliest time at which the tree has five such clades, and pool the smaller clades. We lay out the tissue as in the TME layout of scCube [9]. The tissue has nine oval tumor nests in a stroma of a first non-tumor cell type, three aggregates of a second type, and a third type that makes up 15% of the stroma. We place the three types following the placement of fibroblasts, T cells and endothelial cells in scCube. Each nest contains 90% tumor cells, and the rest of each nest consists of cells of the second and third types. The three non-tumor types are synthetic and differ only by a random shift of their gene expression. We assign the tumor sites of this layout to the tumor cells along the tree. We cut the sites of each clade into one contiguous portion per child clade with the size of the child clade, starting at the root. We make each cut along the principal axis of the sites of the clade after turning it by a random angle with a standard deviation of one radian. We fill the other sites with the three non-tumor cell types. The tissue contains about one cell per 100 $\mu m^2$ (Figure S2). We model the gene expression of each cell over 2,000 genes as the sum of a baseline, a cell type program, heritable programs that evolve along the true tree as Brownian motion, and positional programs of strength 0.1 given by random Fourier features of the cell locations. We simulate the counts from Poisson distributions with an expected library size of 1,000. We split the cells at random in two halves and measure one half as by Visium with spots of 55 $\mu$m on a hexagonal lattice 100 $\mu$m apart. Each spot contains the cells whose centers are within it (12 on average), and we discard the cells between spots. A quarter of the other half (about 4,000 cells) forms the dissociated sample, and we read the lineage of 70% of its tumor cells.

We vary one property of the tumor in each scenario: the strength of the heritable programs (0.03, 0.06, 0.1 and 0.3), the number of clones whose cells are dispersed over the section (none, one or two), or the number of clones (three, five or ten). We keep the other properties at the base setting of heritable programs of strength 0.06, no dispersed clone and five clones. To disperse a clone, we exchange the positions of its cells with those of random tumor cells of the clones that are not dispersed. We simulate each setting with two seeds. We chose $\alpha = 0.2$ and $\beta = 1/2$ for SpOTr on a third seed of the base setting, which the benchmark does not score, and used these settings throughout. They had the highest mean per-clone Pearson correlation over a grid of $\alpha$ from 0 to 0.4 and $\beta$ from 1/4 to 3/4.

##### Scores

We score a mapping per clone as in [10]. We compare the true proportion of the clone among the cancer cells with a clone in each spot with its estimated proportion on all spots that contain at least one such cell. We count a spot with no mapped cells as zero for every clone. We compute the Pearson correlation on these proportions, the root mean squared error on the standardized proportions, the structural similarity on the proportions divided by their maximum, and the Jensen–Shannon divergence on the proportions divided by their sum. We average each score over all clones with at least 20 true cells, which include the pooled smaller clades.

##### Real data

*PEtracer.* We use the time-calibrated tree, clades and cells of [11]. We split the cells of section M3-4 at random into two halves, build synthetic Visium spots from one half as in the simulations, and draw 8,000 cells at random from the other half as the dissociated sample. We score the mappings as in the simulations.

*HTAN.* Case HT260C1 of [12] is a colorectal liver metastasis with single-nucleus multiome data and a Visium section from the same block (HT260C1-Th1K1Fc2U1). We split the genes measured in both at random in two halves. All methods fit genes of the first half that pass the gene filter of cell2location. We compute the copy number aberration (CNA) signal of the spots, which serves as a reference, from the second half only. We call clones A and B, the sub-lineages B1 and B2 of clone B, and the sub-lineages B1a and B1b of B1 from the CNA scores of the nuclei, and build the tree from these CNA scores (Clones and tree of HT260C1, below). We do not give B1a and B1b to any method.

##### Clones and tree of HT260C1

We compute a CNA score for each chromosome arm of each nucleus from gene expression as in inferCNV [13]. We use the genes detected in at least 5% of the nuclei, normalize the counts of each nucleus to 10,000, log-transform them, and subtract from each gene its mean in the non-tumor nuclei. We then smooth these values along each chromosome with a moving average over 101 genes, subtract the median over the genes of each nucleus, and average the genes of each chromosome arm. We also compute a CNA score for each chromosome arm of each nucleus from chromatin accessibility as the log2 ratio between the fraction of the peak counts of the nucleus on the chromosome arm and the mean of this fraction in the non-tumor nuclei.

Clustering the CNA scores of all tumor nuclei directly would group them partly by sequencing depth, since the CNA scores of shallow nuclei are noisier and shift with depth. We therefore define the clones on the nuclei with the most counts and call each nucleus from a likelihood ratio on the counts of the nucleus, which accounts for differences in sequencing depth. To call the clones, we first regress the log total counts out of the CNA scores of the tumor nuclei. We then standardize each chromosome arm, split the 60% of tumor nuclei with the most counts into two groups by Ward clustering of their CNA scores, and select the chromosome arms whose CNA scores differ between the two groups by at least 0.5 standard deviations and on which most genes change in the same direction. We then compute the likelihood ratio of the counts of each of these nuclei on the genes of the selected chromosome arms under the two groups. The likelihood is multinomial, and we estimate the proportions of counts on each chromosome arm in each group from its nuclei among the 40% with the most counts. We call a nucleus A or B when the posterior probability of either clone reaches 0.9 under equal priors. We split clone B into B1 and B2 with a Gaussian mixture model (GMM) with two components along the first principal component of the CNA scores of its nuclei. Then, we call the remaining tumor nuclei by adding the evidence of their chromatin accessibility to that of their expression. We train a logistic regression of the clone (A or B) on the chromatin accessibility CNA scores of the nuclei called so far, after regressing out the log total peak counts. The log-odds of B against A of each remaining nucleus is the log-likelihood ratio of the counts of the nucleus plus the log-odds that the regression predicts minus the log-odds of B among the nuclei the regression is trained on, so that we count the prior only once. We call the nucleus B when the resulting posterior probability is at least 0.9 and call it A when the posterior probability is at most 0.1. We call B1 and B2 among the nuclei of B in the same way but use the log-odds of the posterior of the GMM in place of the log-likelihood ratio. We leave out the tumor nuclei that remain uncalled and the tumor nuclei in expression clusters (Leiden) of which at least 30% are non-tumor nuclei. We split B1 into the sub-lineages B1a and B1b with a GMM with two components along the first principal component of the CNA scores of its nuclei. We compute these CNA scores without the smoothing and the subtraction of the median and regress out the log total counts. We build the tree from the CNA scores of the tumor nuclei after regressing out the log total counts and standardizing each chromosome arm. We cluster the nuclei of each of the clones A, B1 and B2 by Ward clustering of their distances and join the three clones by the mean distances between their nuclei. We fit the GMMs and the L2-penalized logistic regressions ($C = 0.1$) with scikit-learn and the Ward clustering with SciPy. We compute the Leiden clusters with scanpy on 2,000 highly variable genes with 30 principal components, 15 neighbors and a resolution of 1.

We use the tumor purity that the authors report for each spot, which they estimated with ESTIMATE [14] from the expression of stromal and immune genes. We keep the spots in the tissue that have a purity estimate and at least 200 counts on the genes that the methods fit. We compute the CNA scores of the spots as for the nuclei from the second half of the genes and use the 20% of spots with the lowest purity as the reference in place of the non-tumor nuclei. We then regress the purity out of the CNA scores of the spots so that they do not vary with the tumor content of the spots, and we standardize each chromosome arm. The CNA signal of a spot for two groups of nuclei is the projection of its CNA scores on the direction across chromosome arms that separates the two groups in the nuclei. We compute the signal for B against A on the spots with at least the median purity, the signal for B1 against B2 on the spots with a positive signal for B, and the signal for B1b against B1a on the territory of B1. The territory of B1 consists of the spots with a signal for B above 0.5 standard deviations and a positive signal for B1. We then score each mapping by its balanced accuracy. We take the proportion of the first group (B, B1 or B1b) in the mapped mass of the two groups on the spots with a signal above 0.5 standard deviations and the proportion of the second group on the spots with a signal below $-0.5$ standard deviations, and we average the two proportions.

### Supplementary Figures

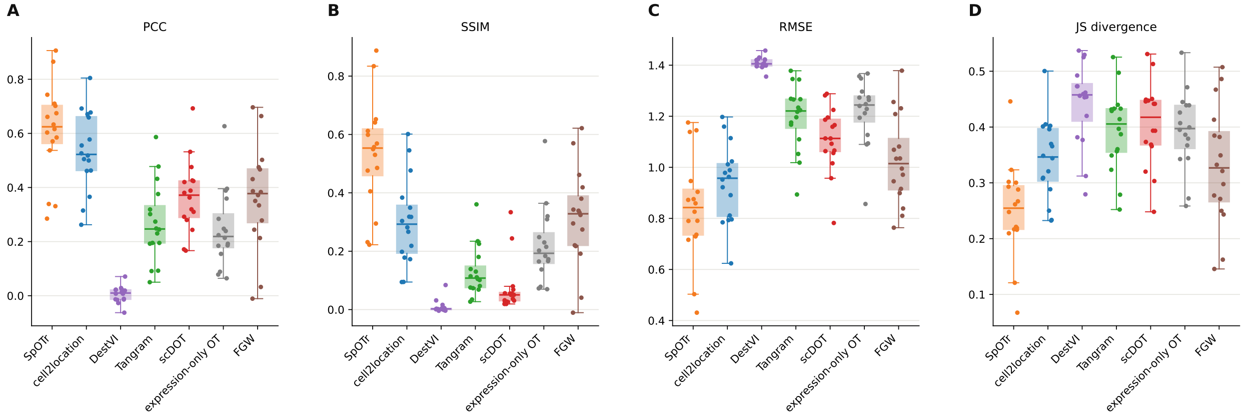


Figure S1: Benchmark metrics on simulated data. We compare the true and the estimated clone proportions of the spots by (A) the Pearson correlation and (B) the structural similarity, for which higher is better, and by (C) the root mean squared error and (D) the Jensen–Shannon divergence, for which lower is better. The points are the simulated tumors of all scenarios.

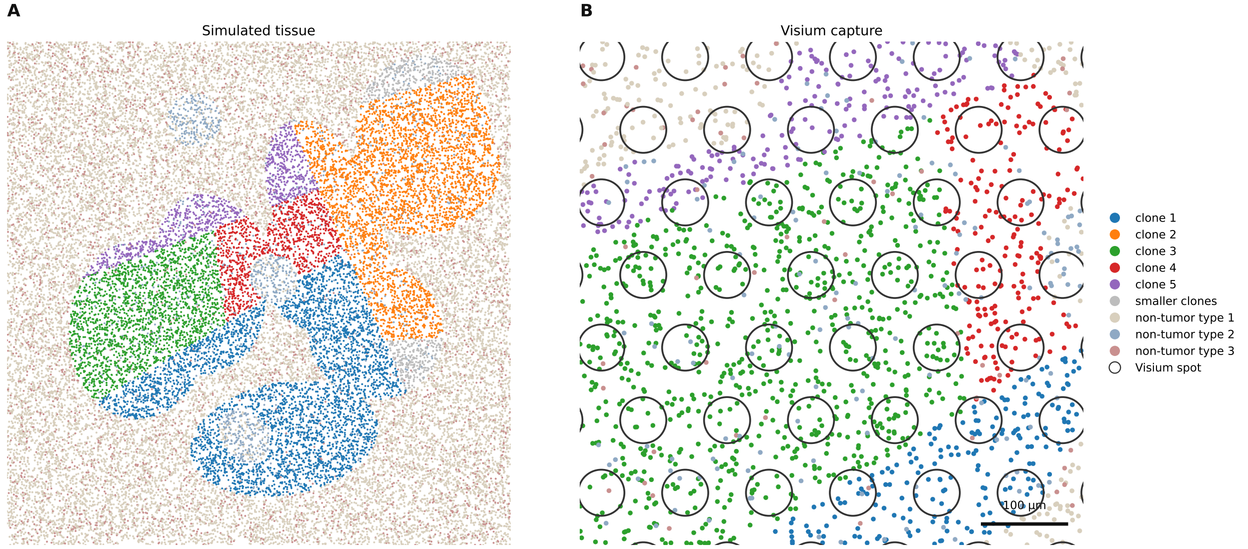


Figure S2: A simulated tumor. (A) All cells of one simulated section, colored by clone for the cancer cells and by type for the three synthetic non-tumor cell types. Each clone has its own territory within the tumor nests of the scCube layout. (B) A window of the measured half of the section with the Visium spots of 55 $\mu$m drawn over its cells. Each spot contains the cells whose centers are within it.

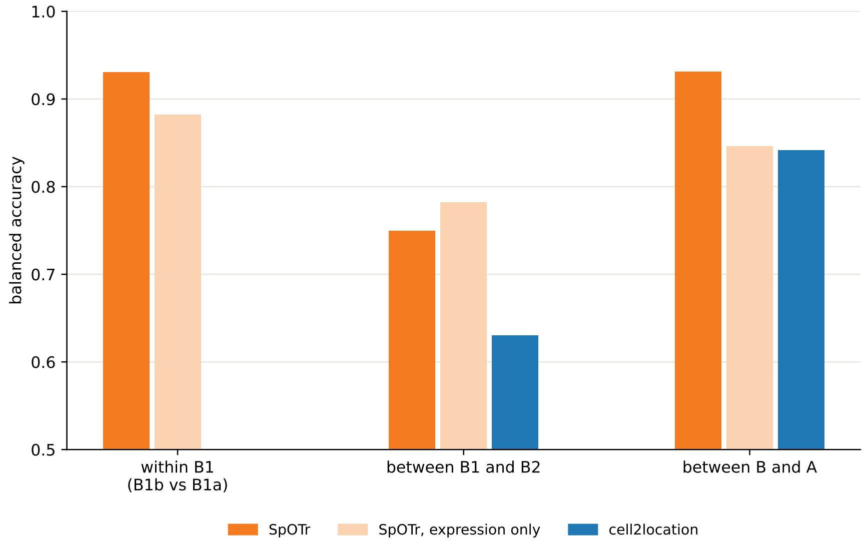


Figure S3: Balanced accuracy on HT260C1. We show the balanced accuracy of the mappings of SpOTr, of SpOTr with expression only and of cell2location against the CNA signal of genes that no method uses, for B1b against B1a within clone B1, for B1 against B2 and for B against A. We do not give B1a and B1b to cell2location.

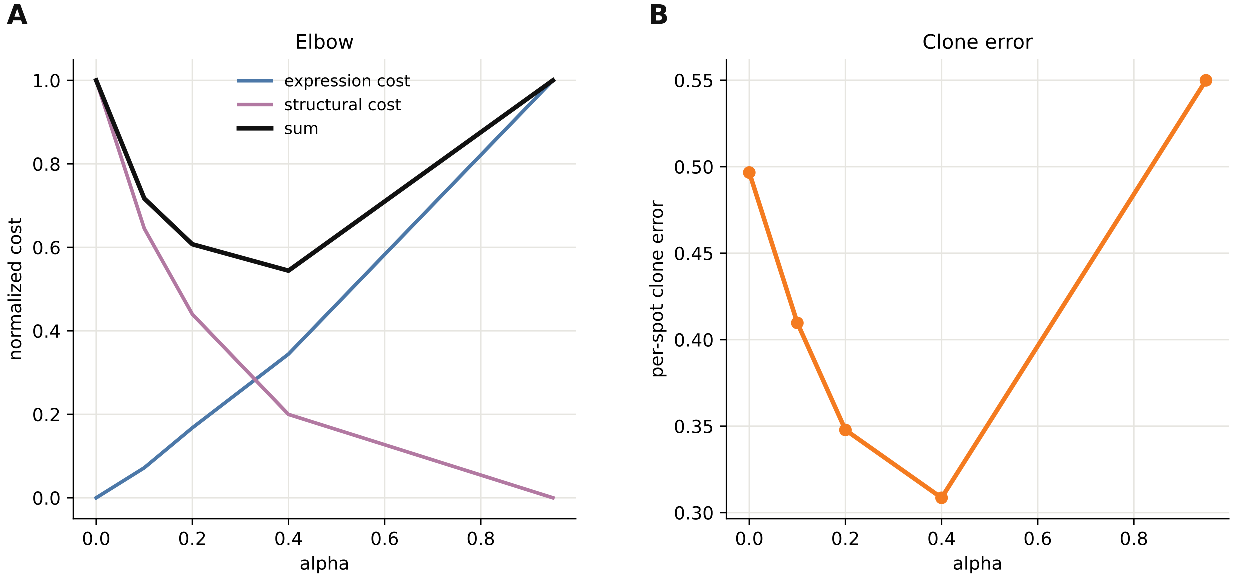


Figure S4: Choosing the structural importance $\alpha$ at the elbow on the simulated tumor of Figure 2. (A) The expression cost and the structural cost of the mapping of SpOTr with the lineage tree and their sum. We normalize each cost between $\alpha = 0$ and $\alpha = 0.95$, and the minimum of the sum is the elbow at $\alpha = 0.4$. (B) The per-spot clone error of each mapping.

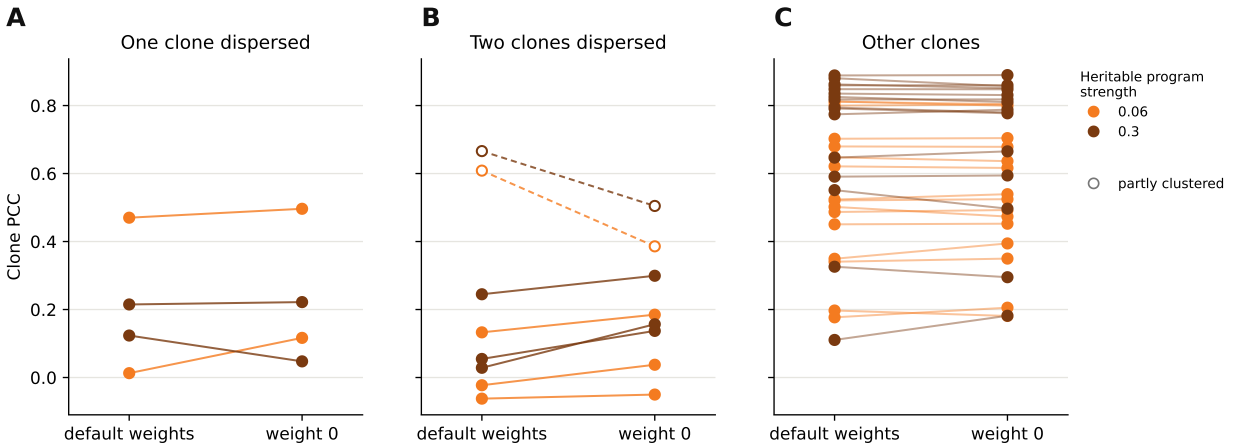


Figure S5: Clade weights on simulated tumors with dispersed clones. We show the per-clone Pearson correlation with the truth of SpOTr with the default clade weights and with weight 0 on the dispersed clones, and we join the two values of each clone by a line. The heritable programs of the clones have strength 0.06 as in the benchmark (orange) or 0.3 (brown). (A) The dispersed clone of the tumors with one dispersed clone. (B) The dispersed clones of the tumors with two dispersed clones. The open markers show a clone that remained partly clustered after the dispersal. (C) The other clones of the same tumors, whose weights we do not change.

#### Supplementary Note 1: the linearized structural term

We derive here the linearized structural term of SpOTr, which we use at every update of the mapping (Methods). The derivation follows the exposition of Gromov–Wasserstein optimal transport in [15, Sec. 10.6.3], and extends it to pair weights between cells and to the capacities of the spots.

##### Notation

Let $n, n' \in \{1, \ldots, N\}$ index the cancer cells and $m, m' \in \{1, \ldots, M\}$ the spots, with $\ell_m \in \mathbb{R}^2$ the position of spot $m$. The rows of the mapping $\gamma \in \mathbb{R}_+^{N \times M}$ sum to one, so that each cancer cell is mapped exactly once, and its columns sum to the capacities of the spots,

$$\sum_m \gamma_{nm} = a_n = 1, \qquad \sum_n \gamma_{nm} = b_m, \tag{1}$$

with row marginals $\mathbf{a} = \gamma \mathbf{1}$ and column marginals $\mathbf{b} = \gamma^\top \mathbf{1}$, where $b_m$ is the capacity of spot $m$ (Methods). $C^T \in \mathbb{R}^{N \times N}$ contains the distances between the cells in the tree, which we scale to the mean distance between the spots. $C^S \in \mathbb{R}^{M \times M}$ contains the distances between the spots, $C^S_{mm'} = \|\ell_m - \ell_{m'}\|$, and $\Omega \in \mathbb{R}^{N \times N}$ contains the weights of the pairs of cells. All three matrices are symmetric. With a tree, $\Omega_{nn'} = 1$ for every pair of distinct cells. With clone labels, $\Omega_{nn'} = \mathbf{1}[k_n = k_{n'}]$ for distinct cells $n$ and $n'$ and $C^T = 0$. We leave out the pairs of a cell with itself, $\Omega_{nn} = 0$. Weights for each clade multiply the pair weights, $\Omega_{nn'} \leftarrow \omega_{k_n} \omega_{k_{n'}} \Omega_{nn'}$. We write $A^{\circ 2}$ for the elementwise square of a matrix $A$ and $\odot$ for the elementwise product.

##### The structural term

The structural term is the weighted squared difference between the tree distances of pairs of cells and the distances between the spots they are mapped to,

$$S(\gamma) = \sum_{n,n',m,m'} \Omega_{nn'} \big(C^T_{nn'} - C^S_{mm'}\big)^2 \gamma_{nm}\, \gamma_{n'm'}. \tag{2}$$

With $\Omega_{nn'} = 1$ for every pair, including those of a cell with itself, this is the Gromov–Wasserstein objective [16]. It is quadratic in $\gamma$ and not convex, and we therefore linearize it at the current mapping and solve the resulting linear problem with entropic regularization [17, 18].

##### Linearization

Expanding the square splits the structural term into three terms,

$$S(\gamma) = \underbrace{\sum_{n,n',m,m'} \Omega_{nn'} (C^T_{nn'})^2 \gamma_{nm}\gamma_{n'm'}}_{\text{A}} + \underbrace{\sum_{n,n',m,m'} \Omega_{nn'} (C^S_{mm'})^2 \gamma_{nm}\gamma_{n'm'}}_{\text{B}} \underbrace{-2 \sum_{n,n',m,m'} \Omega_{nn'}\, C^T_{nn'}\, C^S_{mm'}\, \gamma_{nm}\gamma_{n'm'}}_{\text{C}}. \tag{3}$$

**A**: Summing over the spots first,

$$\text{A} = \sum_{n,n'} \Omega_{nn'} (C^T_{nn'})^2 \sum_{m,m'} \gamma_{nm}\gamma_{n'm'} = \sum_{n,n'} \Omega_{nn'} (C^T_{nn'})^2\, a_n a_{n'}, \tag{4}$$

and since $\partial a_n / \partial \gamma_{nm} = 1$ and $\Omega$ and $C^T$ are symmetric,

$$\frac{\partial \text{A}}{\partial \gamma_{nm}} = 2 \sum_{n'} \Omega_{nn'} (C^T_{nn'})^2\, a_{n'} = 2\big[(\Omega \odot (C^T)^{\circ 2})\, \mathbf{a}\big]_n. \tag{5}$$

**B**: Summing over the cells first,

$$\text{B} = \sum_{m,m'} (C^S_{mm'})^2 \sum_{n,n'} \Omega_{nn'}\, \gamma_{nm}\gamma_{n'm'}. \tag{6}$$

Without the weights, the inner sum is $b_m b_{m'}$, which is set by the capacities, and B is then a constant. With the weights, the inner sum includes only the weighted pairs of cells, and it depends on which cells are mapped to which spots. Since $\Omega$ and $C^S$ are symmetric,

$$\frac{\partial \text{B}}{\partial \gamma_{nm}} = 2 \sum_{m'} (C^S_{mm'})^2 \sum_{n'} \Omega_{nn'}\, \gamma_{n'm'} = 2\big[\Omega\, \gamma\, (C^S)^{\circ 2}\big]_{nm}. \tag{7}$$

**C**: Likewise,

$$\frac{\partial \mathrm{C}}{\partial \gamma_{nm}} = 2 \sum_{n',m'} \Omega_{nn'}\, C^T_{nn'}\, \gamma_{n'm'}\, C^S_{m'm} = 2\big[(\Omega \odot C^T)\, \gamma\, C^S\big]_{nm}. \tag{8}$$

Adding the three derivatives gives the gradient $2L^{\Omega}$ of the structural term, with

$$L^{\Omega}_{nm} = \big[(\Omega \odot (C^T)^{\circ 2})\, \mathbf{a}\big]_n + \big[\Omega\, \gamma\, (C^S)^{\circ 2}\big]_{nm} - 2\, \big[(\Omega \odot C^T)\, \gamma\, C^S\big]_{nm}, \tag{9}$$

where $L^{\Omega}_{nm}$ is the cost of placing cell $n$ in spot $m$ given the current mapping of the other cells. We absorb the factor of 2 in the strength of the term, which we calibrate against the likelihood of the spots (Methods). With $\Omega_{nn'} = 1$ for every pair, $\Omega\gamma = \mathbf{1}\mathbf{b}^\top$, and eq. (9) becomes the linearized cost of Gromov–Wasserstein [16],

$$L_{nm} = \big[(C^T)^{\circ 2}\, \mathbf{a}\big]_n + \big[(C^S)^{\circ 2}\, \mathbf{b}\big]_m - 2\, \big[C^T \gamma\, C^S\big]_{nm}. \tag{10}$$

With clone labels, $C^T = 0$ and only the second term remains. This term is the sum of the squared distances between spot $m$ and the spots to which the other cells of the clone of cell $n$ are mapped. The term is thus small when the cells of each clone are mapped close together, and it does not include pairs of cells from different clones.

##### Computing the term

The second term does not require the matrix of squared distances between the spots. We write $G = \Omega\, \gamma$. Since $(C^S_{mm'})^2 = \|\ell_m\|^2 + \|\ell_{m'}\|^2 - 2\, \ell_m^\top \ell_{m'}$, we have

$$\big[G\, (C^S)^{\circ 2}\big]_{nm} = \|\ell_m\|^2 \sum_{m'} G_{nm'} + \sum_{m'} G_{nm'}\, \|\ell_{m'}\|^2 - 2\, \ell_m^\top \sum_{m'} G_{nm'}\, \ell_{m'}, \tag{11}$$

and computing this costs as much as the mapping itself once $G$ is known. With a tree, $G = \mathbf{1}\mathbf{b}^\top - \gamma$. With clone labels, $\Omega$ is block diagonal, and $G$ is the sum of the mapping over the other cells of each clone. Only the third term, which is needed for a tree, multiplies the mapping by the matrix of distances between the spots.

##### Entropic optimal transport with the linearized cost

We linearize the whole objective at the current mapping at every update, and obtain the cost of placing cell $n$ in spot $m$ as

$$\Lambda_{nm} = -\frac{\partial \log p(Y \mid \theta, \gamma)}{\partial \gamma_{nm}} + \beta' C^F_{nm} + \alpha' L^{\Omega}_{nm}, \tag{12}$$

where $\beta'$ and $\alpha'$ are the calibrated strengths of the expression cost and of the structural term (Methods). The entropic optimal transport problem with this cost,

$$\arg\min_{\gamma} \sum_{n,m} \gamma_{nm} \Lambda_{nm} + \gamma_{nm} \log \gamma_{nm} \quad \text{subject to} \quad \gamma \mathbf{1} = \mathbf{1}, \quad \gamma^\top \mathbf{1} = \mathbf{b}, \tag{13}$$

has the solution $\gamma_{nm} = \exp(f_n + g_m - \Lambda_{nm})$, whose potentials we find by alternating the Sinkhorn updates in the log domain [15, 18],

$$f_n \leftarrow -\log \sum_m \exp(g_m - \Lambda_{nm}), \qquad g_m \leftarrow \log b_m - \log \sum_n \exp(f_n - \Lambda_{nm}), \tag{14}$$

until both marginals are met. We then move the mapping towards this solution by a step chosen on the exact objective, and repeat the linearization at the new mapping (Methods).

[6] Titouan Vayer, Laetita Chapel, Rémi Flamary, Romain Tavenard, and Nicolas Courty. Fused gromov-wasserstein distance for structured objects: theoretical foundations and mathematical properties. *arXiv preprint arXiv:1811.02834*, 2018.

[13] Trinity CTAT Project. inferCNV of the Trinity CTAT Project. `https://github.com/broadinstitute/inferCNV`.

[14] Kosuke Yoshihara, Maria Shahmoradgoli, Emmanuel Martínez, Rahulsimham Vegesna, Hoon Kim, Wandaliz Torres-Garcia, Victor Treviño, Hui Shen, Peter W Laird, Douglas A Levine, et al. Inferring tumour purity and stromal and immune cell admixture from expression data. *Nature Communications*, 4:2612, 2013.

[15] Gabriel Peyré and Marco Cuturi. Computational Optimal Transport, March 2020. arXiv:1803.00567 [stat].

[16] Gabriel Peyré, Marco Cuturi, and Justin Solomon. Gromov–Wasserstein averaging of kernel and distance matrices. In *Proceedings of the 33rd International Conference on Machine Learning*, pages 2664–2672, 2016.

[17] Justin Solomon, Gabriel Peyré, Vladimir G Kim, and Suvrit Sra. Entropic metric alignment for correspondence problems. *ACM Transactions on Graphics (ToG)*, 35(4):1–13, 2016.

[18] Marco Cuturi. Sinkhorn Distances: Lightspeed Computation of Optimal Transportation Distances, June 2013. arXiv:1306.0895 [stat].
